# Flowering cues in a Costa Rican cloud forest: analyzing the effect of climate

**DOI:** 10.1101/2022.07.20.500362

**Authors:** Steven E. Travers, Ned A. Dochtermann

## Abstract

The influence of a changing climate on the phenology of organisms in a region is dependent on how regional climate cues or modifies the timing of local life history events and how those cues are changing over time. There is extensive evidence of phenolological shifts in flowering time over the past 50 years in response to increasing temperatures in temperate regions, but far less is known about tropical regions where seasonality is less temperature driven. We examined historical datasets of flowering patterns in two guilds of ornithophilous plants in the montane cloud forests of Monteverde, Costa Rica in order to identify environmental cues for flowering in nine species of plant that are important resources for hummingbirds. Bimonthly censuses of flower production were used to quantify flower production during two sampling periods:1981-1983, 1986-1991., the species studied here appear to cue flowering patterns to either accumulated drought units or a combination of accumulated drought units and chill units prior to flowering. These results have implications for how tropical cloud forest plants will respond to climate change to the extent that drought and chill patterns are changing with time.

## 1. INTRODUCTION

One of the best documented impacts of global climate change on organisms is a shift in the timing of flowering of plant species over the past fifty years. Flowering phenology, the timing of flower production by seasonally reproducing plants has shifted earlier in a wide variety of species (Parmesan 2007, Wolkovich et al. 2012, CaraDonna et al. 2014). In particular, there is strong evidence that temperate species that flower in the spring have been shifting flowering earlier since the 1970s, when global annual temperatures began to increase consistently (Miller-Rushing and Primack 2008, Fitter and Fitter 2002, Menzel et al. 2006). The species deviating most from historical patterns in temperate latitudes are the species flowering earliest in the spring (Wolkovich et al. 2012, Theobald et al. 2017). Because flowering phenology can play an important role in plant reproductive success, the historical timing of flowering is believed to have evolved according to adaptive peaks (Rathke and Lacy 1985). Climate change related shifts in phenology may disrupt important symbiotic relationships such as between plants and pollinators by decreasing synchrony in phenological patterns that have evolved over long periods of time; a consequence that may play an important role in the ecology and evolution of plant species(Kharouba et al. 2018, Post et al. 2008, Thackeray et al. 2016). Indeed, recent evidence suggests that phenological sensitivity to climate change may in fact be linked to individual fitness and long-term species persistence (Primack et al. 2009, Cleland et al 2012).

It is clear that many spring flowering species have changed their phenology as global CO_2_ levels and temperatures have increased and regional climates have changed. However, it is less clear how broadly these patterns apply across all latitudes. The majority of evidence for phenological shifts in flowering is based on studies of plant populations at temperate latitudes (Cook et al. 2012). However, there are reasons to suspect that tropical plant populations might respond differently to climate change (Abernathy et al. 2018). Because growing seasons in temperate latitudes are often restricted by sub-freezing temperatures and thus are increasing in length with increasing global temperatures, temperate plants may now experience temperature cues for flowering earlier than they did historically. In contrast, plants at tropical latitudes are often capable of growing throughout the year because of the lack of winter and frost. In contrast to temperate latitudes, the growing seasons may be dependent on precipitation patterns (e.g. Augsberger 1981) or other environmental cues. Thus, flowering patterns in tropical plants may track shifts in non-temperature environmental variables that may have also changed over the past fifty years. There have been relatively few studies of the effects of climate change on flowering phenology in tropical ecosystems (Morellato et al. 2013).

What do we know about long-term phenology patterns in tropical communities? In general, analyses of the few long-term datasets available describe relationships between either flowering and environmental variables other than air temperature or fruiting and environmental variables other than temperature. The ongoing study of flowering phenology in a lowland moist forest in Panama (Barro Colorado Island) has provided a rich dataset for analyses of both cues for flowering and correlates of flowering intensity in that environment. Seasonal and inter-annual variance in flower production on BCI is best explained by patterns of irradiance and cloudiness, suggesting that cues for initiating flowering in at least some species may come from levels of photosynthetically active radiation (Zimmerman et al. 2007, Wright and Calderon 2018). Pau et al. (2018) found that for the same long-term study the best predictor of decadal patterns in intensity of flowering was global carbon dioxide although correlated changes in irradiance may also play a role. As global CO_2_ levels have increased since the 1970s so has flower production in many of the species in the study. Several studies have analyzed long-term patterns in fruiting phenology which tends to be strongly correlated with flowering phenology (Ettinger et al. 2018). As in flowering phenology, irradiance has been implicated as an important driver of fruiting intensity in trees in tropical Uganda (Chapman et al. 2018). Seasonal differences in rainfall are correlated with fruiting patterns in many tropical ecosystems (Mendoza et al. 2017) and relationships between long-term changes in precipitation and shifts in phenology have been observed in Madagascar (Dunham et al. 2018) and Uganda study sites (Chapman et al. 2005). Perhaps because long-term datasets of tropical plant phenology represent a diverse set of species and ecosystems, no single environmental driver of phenology shifts has emerged despite evidence of year-to-year variance in reproductive intensity and phenology (Polansky and Boesch 2013, Mendoza et al. 2018). These studies indicate that variables such as the abundance and timing of flower and fruit production are shifting in some tropical communities as the global climate changes. However, we are only beginning to understand the environmental drivers and only in a subset of tropical communities.

Understanding the implications of phenological shifts for co-evolved mutualisms requires an understanding of long-term phenological patterns of tropical plant species that are dependent on both other taxa and on environmental correlates. We examined data from 8 years of phenological observations on a collection of cloud forest plants that are dependent on hummingbirds for pollination to better understand how climate change has influenced the timing and abundance of flowering in recent decades.

Neotropical, ornithophilous plant species are ideal for examining the impact of climate change on flowering phenology and, ultimately, on mutualistic interactions. As a result of diffuse coevolution (Feinsinger 1983), the hummingbird-pollinated plants in the cloud forests of Monteverde, Costa Rica have evolved morphological characteristics that separate them loosely into two guilds depending on whether they are pollinated by short-billed or long, curve-billed hummingbirds (Stiles 1981, Feinsinger and Colwell 1978, Feinsinger et al. 1986). The flowering phenology is staggered among these plant species resulting in a continuous supply of nectar with a spike in availability in the transition to dry season (November to February; Feinsinger et al. 1988). However, it is unclear how (or if) these phenological patterns have shifted over the past fifty years as has been seen in temperate latitudes. Tropical montane forests are also one of the least-studied of tropical communities in regard to phenology (Mendoza et al. 2017).

The Monteverde Cloud Forest reserve is located above 1500 m in the Cordillera de Tilarán mountain range of northwestern Costa Rica. Historically, the forest remained wet year-round due to northeasterly trade-winds bringing moisture from the Caribbean sea producing clouds and mist following adiabatic cooling (Clark et al, 2000). In the last 40 years, there have been two climatic trends. First, the daily minimum temperature has increased with increasing global temperature. Second, the number of sequential dry days per year has increased over time (Pounds et al. 1999, 2006). This precipitation shift has closely mirrored the sea surface temperatures of the El Niño Southern Oscillation (ENSO) in the Pacific (Pounds et al. 2006). These long-term shifts may well influence both the drivers of flowering phenology as well as the production of floral resources in neotropical cloud forests. However, as for most tropical biomes long-term phenological data is scarce.

We utilized historical datasets on the phenology of ornithophilous plants from extensive studies on hummingbird-plant associations in the cloud forest reserve of Monteverde. Our study combined these datasets with current observations of flowering phenology of the same species to examine the relationship between changing climate, environmental variability and flower production in one of the richest hummingbird communities in the world (Feinsinger 1977). The goal of our study is to better understand how climate change is impacting tropical flowering patterns by answering the following questions: 1) What are the cues for the timing of flowering and flower abundance in a suite of hummingbird-pollinated plants from the tropical rainforests of Costa Rica? and 2) How do the requirements for flowering initiation compare among different plant species?

## 2. METHODS

### 2.1 Study Plants

We focused on nine plant species whose historical flowering peaks take place during the transitional and dry seasons in the Monteverde cloud forest (January – April). There are many plant species that flower during these months, but we focused on nine species that have traits associated with the hummingbird-pollinated syndrome including tubular corollas, production of copious amounts of nectar with high sucrose content and reddish coloration of the floral parts (Cronk and Ojeda, 2008).

The focal species we chose can be split into associations with one of two hummingbird guilds: short-billed and long-billed. Plants associated with visitation by long-billed hummingbirds are *Razisea spicata* (Acanthaceae), *Centropogon solanifolius* (Campanulaceae), *Columnea magnifica* (Gesneriaceae), *Drymonia rubra* (Gesneriaceae) and *Hillia triflora* (Rubiaceae). Plants associated with visitation by short-billed hummingbirds are *Hansteinia blepharorachis* (Acanthaceae), *Besleria triflora* (Gesneriaceae), *Psychotria elata* (Rubiaceae) and *Palicourea lasiorhachis* (Rubiaceae). Some short-billed hummingbirds can extract nectar from longer flowers by circumventing the floral tube and robbing from nectar spurs although this is not common at Monteverde (Wolfe et al. 1972, Stiles 1977). All of these species have seasonal (temporally dispersed) flowering periods with a single peak that occurs within the dry season (January-April). Epiphytic plants higher in the canopy were not included due to the restrictions of our sampling method.

### 2.2 Flower sampling

The flower census data were collected by Peter Feinsinger and colleagues and later published in a series of important papers describing the ecology and coevolution of hummingbird communities and the plants serving as their nectar sources (Feinsinger et al. 1986, 1988). They censused the flower production of 26 species including the nine focal species in this study over time by counting the number of open flowers observed from the ground at regular temporal intervals. In order to standardize the number of plants censused for open flowers over time, a specified route and distance were always used for replicate censuses over time within a given sampling period (see below for details). At each census date the number of open flowers of a given species were counted by an observer walking the sampling route and scanning each side of the trail (< 2 m deep) and above the trail (< 3 m up). In order to be counted, the flowers needed to be visible without binoculars and open to pollinators based on corolla coloration and presentation (W. Zuchowsi, pers. comm)

There were two distinct sampling periods in this study. The first period (P1) was between July 1981 and June 1983. The second period (P2) was between August 1986 and August 1991. All sampling was conducted within the boundaries of the Monteverde Cloud Forest reserve in Puntarenas Province, Costa Rica. Sampling during periods P1 and P2, as part of two broader studies was conducted at 14 different sites within the reserve between 1500m and 1600 m elevation (Feinsinger et al. 1988, Linhart et al. 1997). Between July 1981 and June 1982 (P1), flowers were counted over the course of one to two days once a month. After June 1982, all sampling intervals were approximately 14 days. In total, flowers were counted in the reserve at 180 different sampling events.

### 2.3 Climate data

Climatological data were collected from multiple sources. Monthly estimates of precipitation were collected near (< 1 km) the Monteverde Reserve (10°17’56” N, 84°48’9” W) by John Campbell from 1956 to 1994 (KNB data archive). Temperature data for 1981 to 1991 were extracted from mean published values that were collected by John Campbell at the same site as the precipitation data (Anchukaitis et al. 2008). We used the program *Engauge* (available online) to determine temperature values from the published figures in Anchukaitis et al. (2008).

### 2.4 Temporal changes in flower abundance

We were interested in determining what environmental factors initiate flowering and potentially regulate the abundance of flowers produced. Our goal was to better understand the relative importance of daylength, precipitation and temperature in determining the monthly quantity of flowers as a correlate of nectar resources. We used a model selection approach to model and predict the relationship between environmental variables and the production of flowers.

Our approach to analyzing the relationship between the three environmental variables and flower production by the focal species in our study is based on modifications of the method used by Chen et al. (2018). Here we assume that the initiation of the flowering period of our focal species is the result of accumulated drought and chilling over a specific time period (n_i_) such that once a critical level of drought and/or chill has accumulated, individual plant species will initiate floral development during a development time period (n_d_) followed by flower production. In the cloud forest of Monteverde, dry periods and cooler temperatures vary widely prior to and during the dry season (Clark et al. 2000). Our modelling approach assumes that seasonal patterns in flowering have evolved in response to historical patterns in these two parameters. In addition, daylength is a cue used by many plant species to initiate flowering (Andrés and Coupland 2012) although daylength varies less over time in the tropics relative to temperate latitudes. In our model, there are two periods - signal accumulation and flower development - that are followed by flowering (see Chen et al.2018). We fit four different combinations of daylength, drought unit accumulation, and chilling unit accumulation to our data in order to examine the relationship between our measures of flowering intensity and the climate prior to floral production (see below).

In order to calculate the accumulated drought units (DU) during the signal accumulation period we used the following equation:

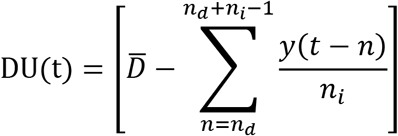

where 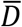 is the threshold level of monthly precipitation, y is monthly precipitation during month t and n_i_ and n_d_ are the numbers of months during the signal accumulation and development periods, respectively. We varied 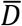 from 100 mm to 400 mm by 20 mm intervals. The length of the signal accumulation period (n_i_ = number of months for signal accumulation) varied from 1 to 6 months and the length of the development period (n_d_ = number of months for flower development) varied from 0 to 3 months both by one-month intervals. All possible combinations of these two time periods were assessed for a total of 504 drought and chill unit combinations.

We calculated chill unit accumulation by using a similar equation that averaged monthly temperatures over the accumulation and development periods:

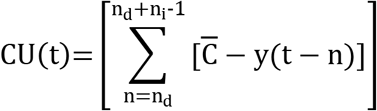

where 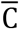 is the threshold temperature, y is the average minimum temperature during month t and n_i_ and n_d_ are the number of months during the signal accumulation and development periods, respectively. Thus, chill units increase with increasing time that includes sufficiently cool periods. The variable 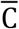 varied from 16 to 19 by intervals of 0.25. The combination of these three variable ranges resulted in a total of 168 chill unit data sets.

Because sampling was conducted at different transects within the reserve between the three sampling periods, it was necessary to standardize flower census counts in order to explore the relationship between precipitation and flowering. We standardized flower counts by first calculating monthly averages for each month in each sample period. We then determined the maximum monthly average value by species and by sampling period. Individual monthly averages were then converted to sampling period specific proportions of the maximum by dividing each monthly average by the relevant maximum. These proportions (F(t)) were used as dependent variables in iterative regressions of F(t) against the independent variables.

The final independent variable tested for effects on flowering was daylength at the time of flowering (DL). The number of minutes of daylight on the 15^th^ of each month was determined from meteorological data collected at Mt. Arenal approximately 19.3 km from the Monteverde reserve. The daylength associated with the flowering month were included as a vector variable in the models as described below. We assumed there was no inter-annual variation in daylength for a particular time period.

We conducted a series of linear regressions on the different combinations of drought units and chill units to examine the relationship between flowering intensity (F(t)) and pre-flowering environment by using each of the following models for comparison:

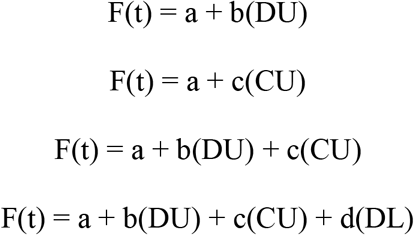

The AIC values (Akaike 1973, 1976) of each fitted linear model were compared and the best fitting model was chosen as the one with the lowest AIC value. We used this approach to train our model to a subset of the complete time series of phenological observations, specifically sample period P2 (1986-1991). The selected model parameters were then tested on the data from sample period one (P1) to determine the ability of the model to predict flowering patterns. We assessed the predictive ability of the models by calculating Pearson correlation coefficients for the relationship between the observed and predicted F(t) values. All analyses were conducted in R 3.4.3 or 3.4.4(R Core Team, 2018).

## 3. RESULTS

### 3.1 Phenological patterns

Each of the nine focal species demonstrated unimodal patterns of flowering (Figure 1) over a 12-month period. All of the species monitored flowered and provided nectar to hummingbirds during the dry season months of January through April (Figure 2). However the start and ending dates of flowering as well as dates of peak flowering varied widely. The species initiating flowering during the dry season include *Besleria triflora, Columnea magnifica, Centropogon solanifolius, Palicourea lasiorachis* and *Psychotria elata*. The other four species, *Razisea spicata, Drymonia rubra, Hillia triflora* and *Hansteinia blepharorachis* start flowering prior to January during the wet season or the transitional season. Peak flowering for the species is spread across the year resulting in a shift in flower availability from species to species over time (Figure 1).

**Figure 1.**
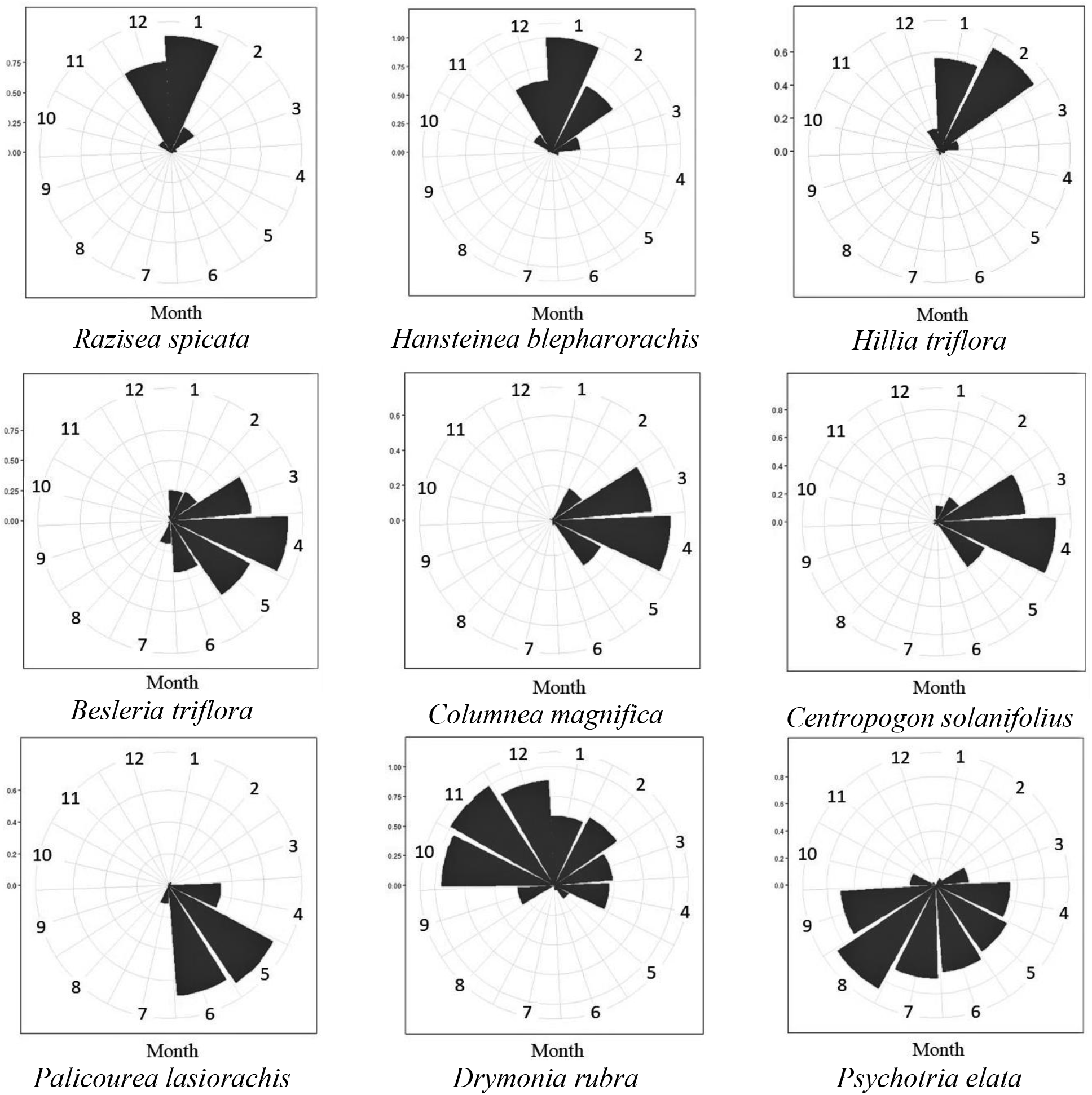
Polar plots of year-round flowering patterns of ten species of hummingbird-pollinated plants in the Monteverde reserve. Monthly values represent an average of the number of flowers open every month during the years 1981-1983 and 1985-1991. Month 1 = January, Month 2 = February, etc.

**Figure 2.**
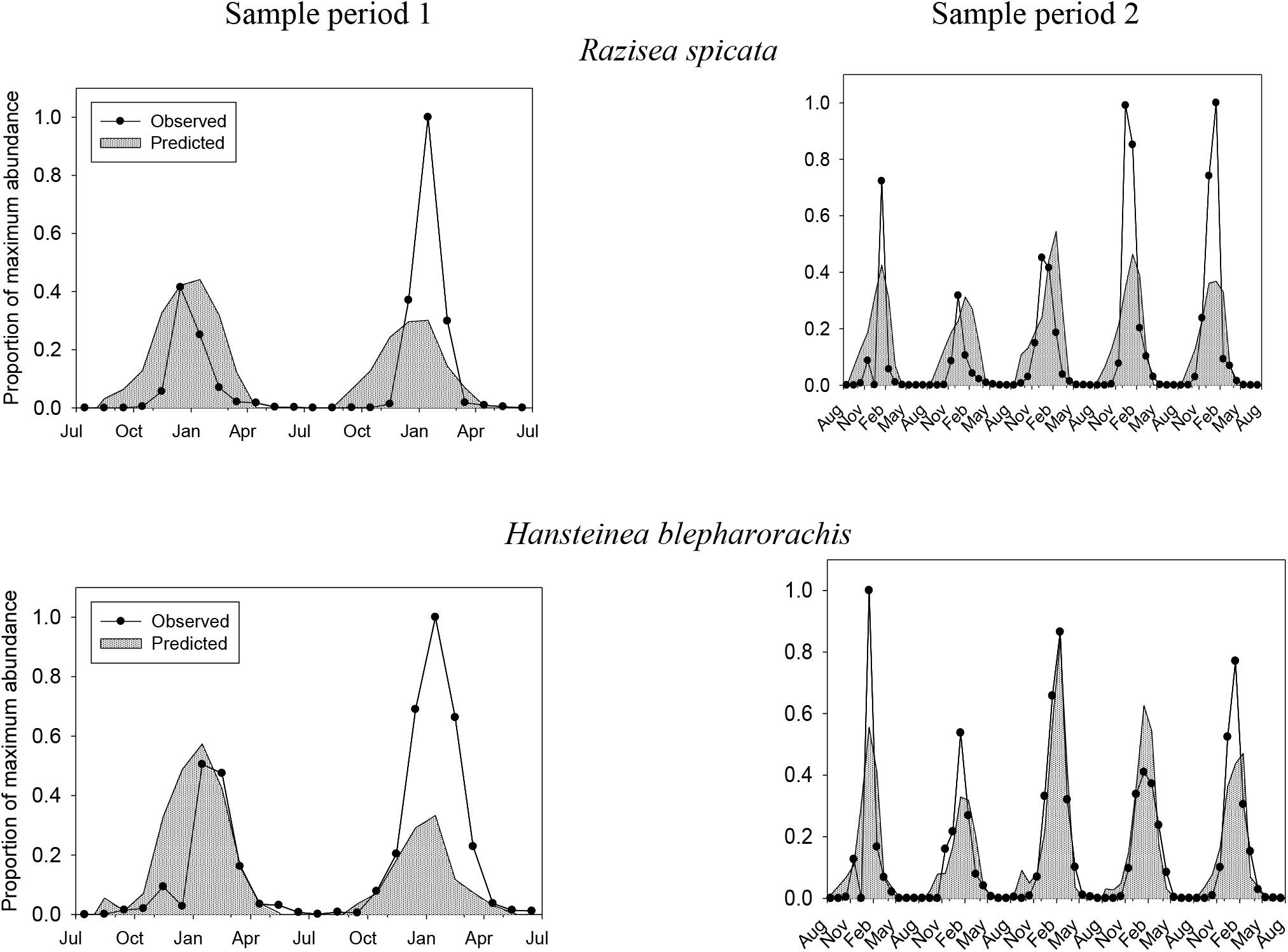
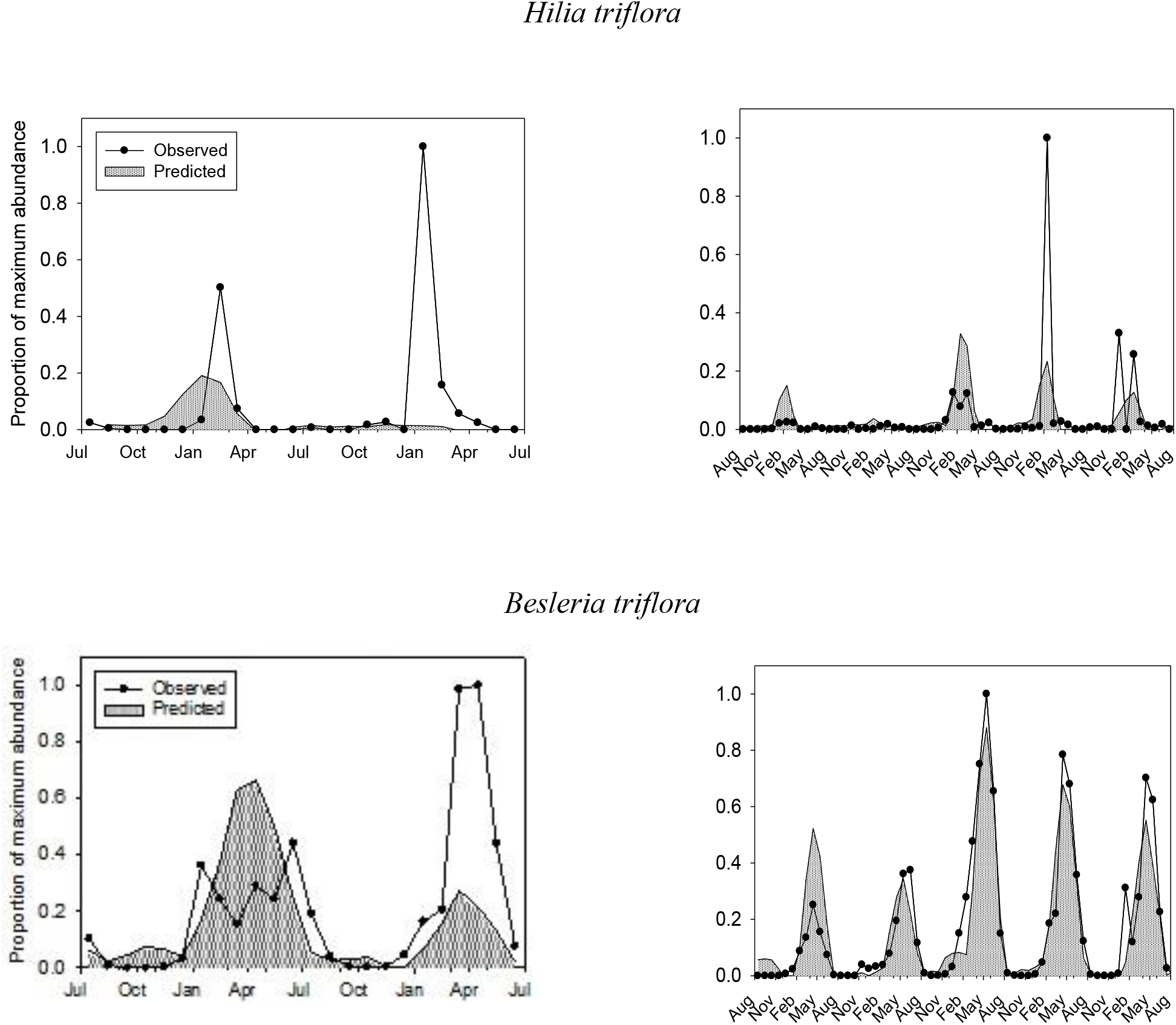
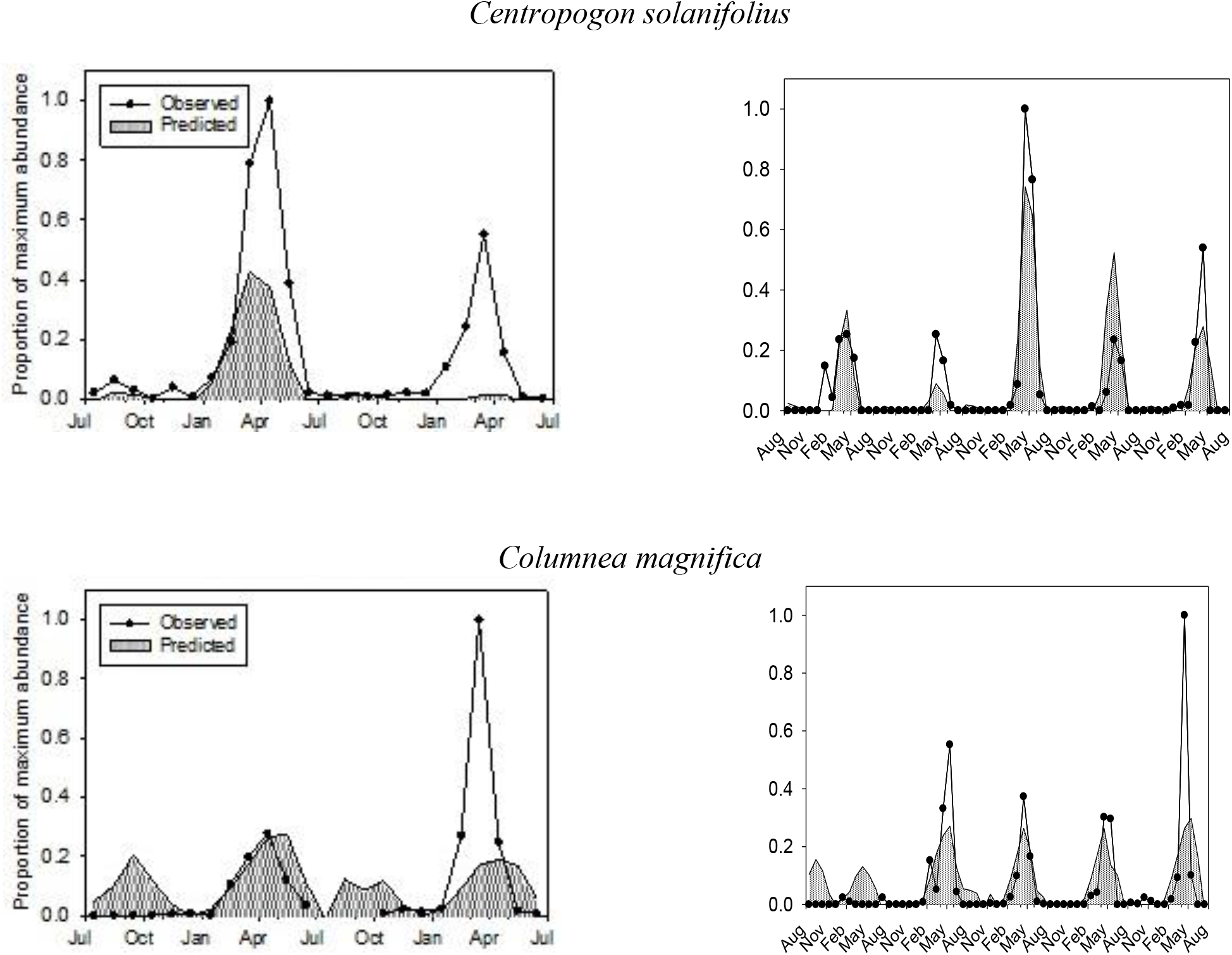
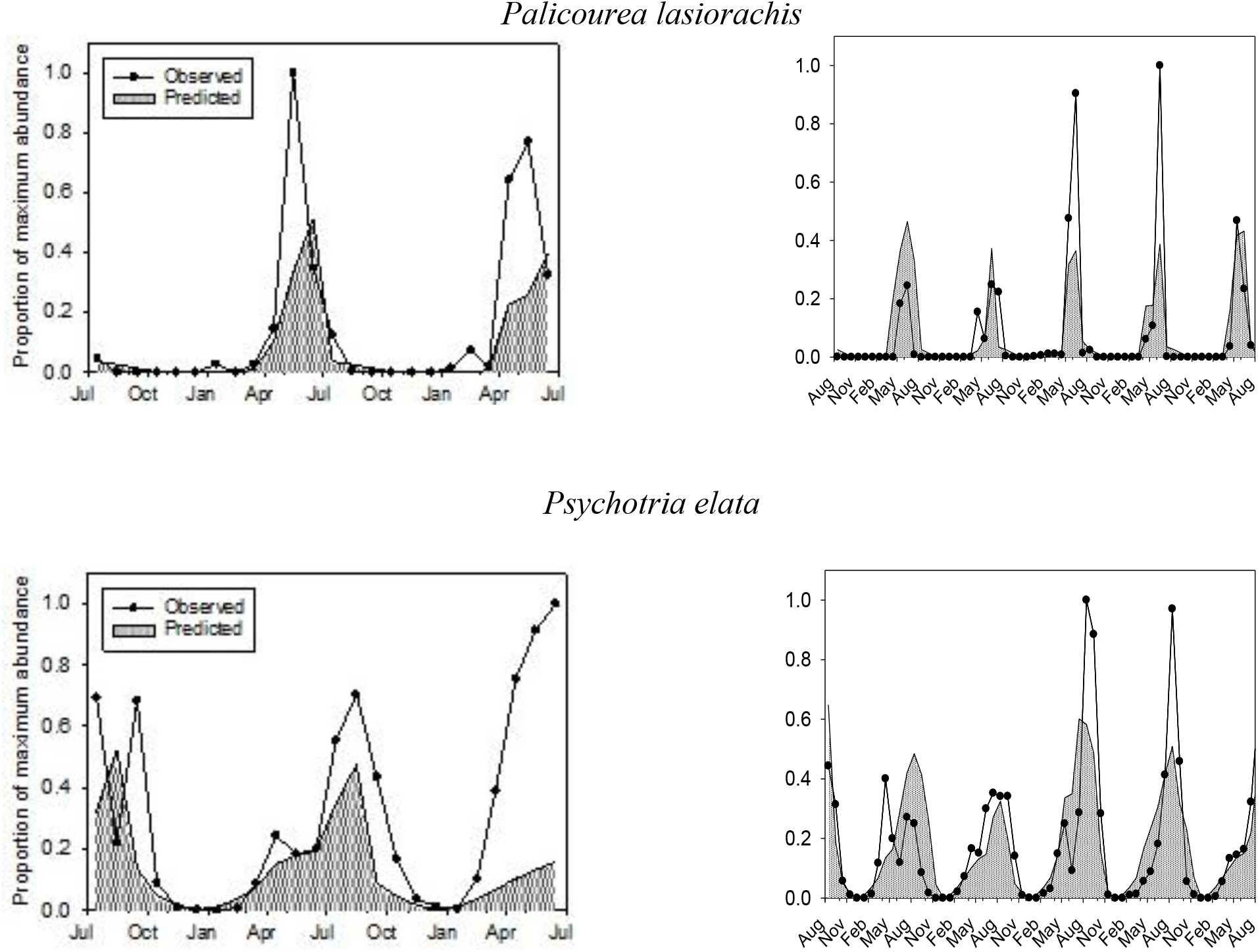
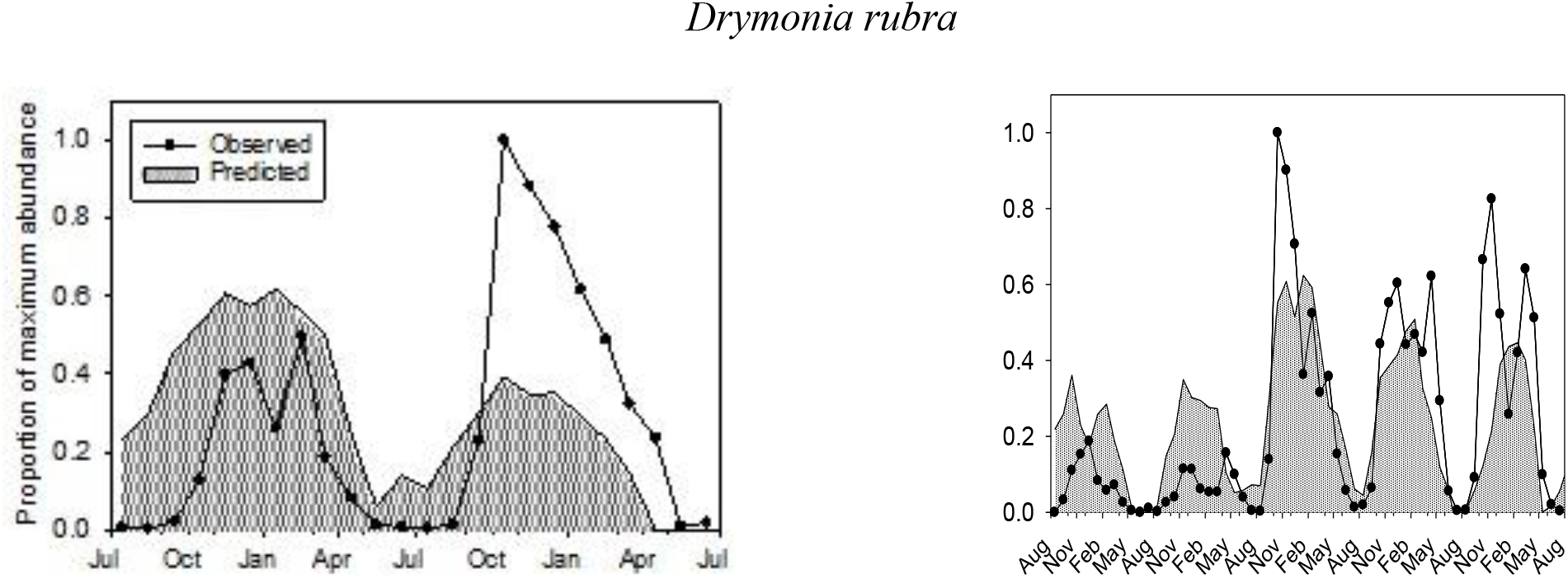
Observed and predicted flowering patterns of focal plant species during three sampling periods (Period 1: 1981-1983, Period 2:1986-1991, Period 3:2017-2018).

### 3.2 Floral abundance patterns of species

The model selection approach to analyzing the relationships between environmental variables and flowering yielded predictive models of flowering for each of the focal species. However, the explanatory ability of selected models varied depending on the sampling period analyzed (Figure 2). As expected, because the different model combinations were fitted to sampling period 2 (1986-1991), the correlation between observed and predicted values were highest for that sampling period. The average correlation coefficient for predicted versus observed values was 0.74 and ranged from 0.5 (*Hilia*) to 0.91(*Centropogon*) for the P2 sampling period. In general, the best models effectively predicted the timing but not necessarily the peak flowering magnitude of each year.

The best-fit models were then used to predict flowering patterns in sampling period one. The models were still predictive though correlation coefficients were lower for the years 1981-1983. The average correlation coefficient for predicted versus observed values was 0.52 and ranged from 0.19 (*Hilia*) to 0.84 (*Centropogon*).

Overall, the best explanatory variable was the drought units prior to flowering compared to either daylength or chill units prior to flowering, suggesting that precipitation patterns are an important regulator of flowering for all focal species. Comparisons of the AIC and delta AIC values for each of the four models fit to P2 data indicate that chill units by themselves explained very little of the variation in flowering (Table 2). Delta AIC values were the largest for a model with chill unit as the single factor relative to the other three models for all nine focal plant species. In contrast, single factor models with drought units produced delta AIC values that were 33% to 100% smaller than for single factor models with chill units. In particular, drought units improved the fit of predictive models over those for chill units extensively in *Palicourea* and *Columnea*.

**Table 2.** The results of model fitting for flower abundance (1986-1991) as a function of different combinations of one to three independent variables (CU=chill units; DU=drought units; and DL=daylength). Best-fit models were chosen on the basis of minimum AIC values; model parameters (*N_i_, N_d_*, 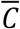 and 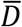) are defined in the text. The overall best models for each species are indicated with bold typeface.

| Species | Models | $N_i$ | $N_d$ | $\bar{C}$ | $\bar{D}$ | AIC | ΔAIC |
| --- | --- | --- | --- | --- | --- | --- | --- |
| *Razisea spicata* | CU | 1 | 0 | 19 | - | 4.86 | 30.8 |
|  | DU | 4 | 1 | - | 380 | -12.5 | 13.5 |
|  | CU+DU | 1 | 0 | 19 | 220 | -19.437 | 6.52 |
|  | **CU+DU+DL** | **1** | **0** | **18.5** | **200** | **-25.957** | **0** |
| *Hansteinea blepharorachis* | CU | 1 | 0 | 16 | - | -2.7 | 81.5 |
|  | DU | 6 | 2 | - | 500 | -36.7 | 47.5 |
|  | CU+DU | 1 | 0 | 18.5 | 340 | -77.8 | 6.4 |
|  | **CU+DU+DL** | **1** | **2** | **18.5** | **480** | **-84.2** | **0** |
| *Hilia triflora* | CU | 1 | 0 | 16 | - | -66.1 | 13.8 |
|  | DU | 2 | 3 | - | 420 | -70.7 | 9.2 |
|  | **CU+DU** | **2** | **0** | **18** | **460** | **-79.9** | **0** |
|  | CU+DU+DL | 2 | 0 | 18 | 460 | -77.9 | 2.0 |
| *Besleria triflora* | CU | 1 | 0 | 16 | - | 3.0 | 101.7 |
|  | DU | 3 | 0 | - | 180 | -47.5 | 51.2 |
|  | **CU+DU** | **3** | **2** | **18.5** | **480** | **-100.1** | **0** |
|  | CU+DU+DL | 3 | 2 | 18.5 | 480 | -98.7 | 0.4 |
| *Centropogon solanifolius* | CU | 1 | 0 | 16 | - | -32.7 | 82.7 |
|  | DU | 3 | 0 | - | 120 | -64.0 | 51.4 |
|  | **CU+DU** | **2** | **2** | **18** | **180** | **-116.9** | **0** |
|  | CU+DU+DL | 2 | 2 | 18 | 180 | -115.4 | 1.5 |
| *Columnea*<br>*magnifica* | CU | 1 | 0 | 16 | - | -45.6 | 25.4 |
|  | DU | 3 | 0 | - | 120 | -69.2 | 1.8 |
|  | CU+DU | 3 | 0 | 16 | 100 | -67.9 | 3.1 |
|  | **CU+DU+DL** | **5** | **2** | **16** | **240** | **-71.0** | **0.0** |
| *Palicourea*<br>*lasiorachis* | CU | 1 | 0 | 16 | - | -24.6 | 40.6 |
|  | **DU** | **3** | **1** | **-** | **100** | **-66.5** | **0** |
|  | CU+DU | 3 | 1 | 16 | 100 | -66.5 | 0 |
|  | CU+DU+DL | 3 | 1 | 16 | 100 | -65.2 | 1.3 |
| *Psychotria elata* | CU | 1 | 0 | 16 | - | -2.0 | 46.6 |
|  | DU | 4 | 3 | - | 220 | -38.1 | 10.5 |
|  | CU+DU | 5 | 3 | 18.5 | 200 | -48.5 | 0.1 |
|  | **CU+DU+DL** | **5** | **3** | **17.5** | **160** | **-48.6** | **0.0** |
| *Drymonia rubra* | CU | 1 | 0 | 16 | - | 13.3 | 32.6 |
|  | DU | 6 | 0 | - | 440 | -13.1 | 6.2 |
|  | **CU+DU** | **5** | **0** | **18.5** | **480** | **-21.3** | **0** |
|  | CU+DU+DL | 5 | 0 | 18.5 | 480 | -19.3 | 2 |

In species other than *Palicourea* and *Columnea*, there was an improvement in the AIC value with the addition of chill units to drought units as independent variables relative to single factor drought unit models. There was a strong interaction between drought unit value and chill unit value in *Centropogon* (Table 2). Ultimately, the best models or models indistinguishable from the best models were those that included all three independent variables. Daylength improved the AIC values the most in *Razisea* and *Hansteinea*.

The model selection approach also identified the most likely signal accumulation or cueing period for flowering. The best fit models invariably had developmental periods (n_d_) that were shorter than the signal accumulation periods (n_i_) which by definition were just prior to the developmental periods. The range of developmental periods was zero to three months while the signal accumulation periods ranged from one to five months (Table 2). The average combination of signal accumulation period and developmental period across species is 4.3 months. The critical 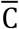 and 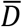 values identified from best fit models varied dramatically among species, even for those with peak flowering similar to one another.

## 4. DISCUSSION

### 4.1 Environmental cues for flowering

The results of our study provide insight into the environmental cues used by cloud forest plants to initiate flowering, Our model selection analysis of flowering abundance indicates that rather than being a function of strictly daylength, flowering is cued by either accumulated drought units or by a combination of chilling and drought (Table 2). Two species stood out as particularly linked to drought units, *Palicourea* and *Columnea*. The other seven species modelled fit better with models incorporating both drought units and chill units. The pattern of flowering in response to a combination of chilling and drought mirrors that found in two other studies of lowland tropical forests where warm temperatures were unimportant relative to these two variables (Chen et al. 2018, Zimmerman et al. 2018). However, Pau et al. (2019) found that leaf and seed phenology in a montane wet forest community tracked either precipitation or temperature patterns, depending on the species, similar to what we found.

Despite reaching peak flowering at different points during the year nearly all of the species seem to flower according to environmental conditions present during the transition from wet season to dry season (Figure 3). Estimation of the signal accumulation period prior to peak flower suggests that signal accumulation most often occured between November and January, regardless of species. From November to January, there is typically a dramatic decrease in daily rainfall and temperatures relative to the rest of year (Clark et al. 2000). These cues bay signal the timing of flowering of the focal species. The few exceptions to the pattern of signal accumulation between November and January include *Hansteinea* which has a signal accumulation period in October. Peak flowering occurs in *Drymonia* much earlier than the other species, it flowers for much longer, and its peak signal accumulation period happens during the beginning of the wet season (Figure 4).

**Figure 3.**
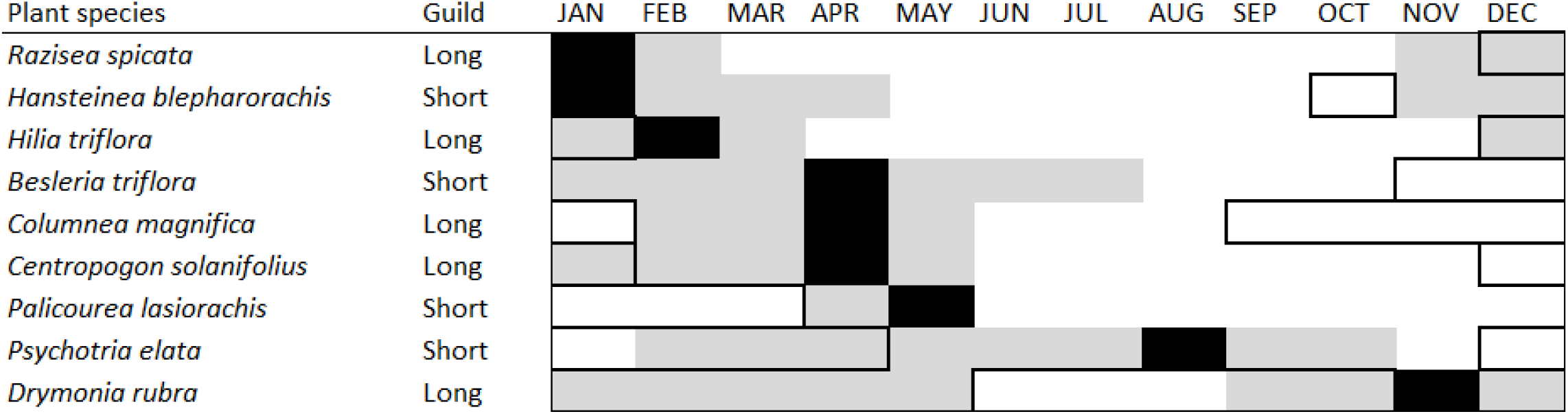
Flowering periods of the focal species studied. Data are based on observation made between 1981 and 1991. Black-fill indicates peak flowering; grey-fill indicates flowering at sub-peak levels; dark-bordered boxes indicate estimated signal accumulation phase prior to peak flowering.

**Figure 4.**
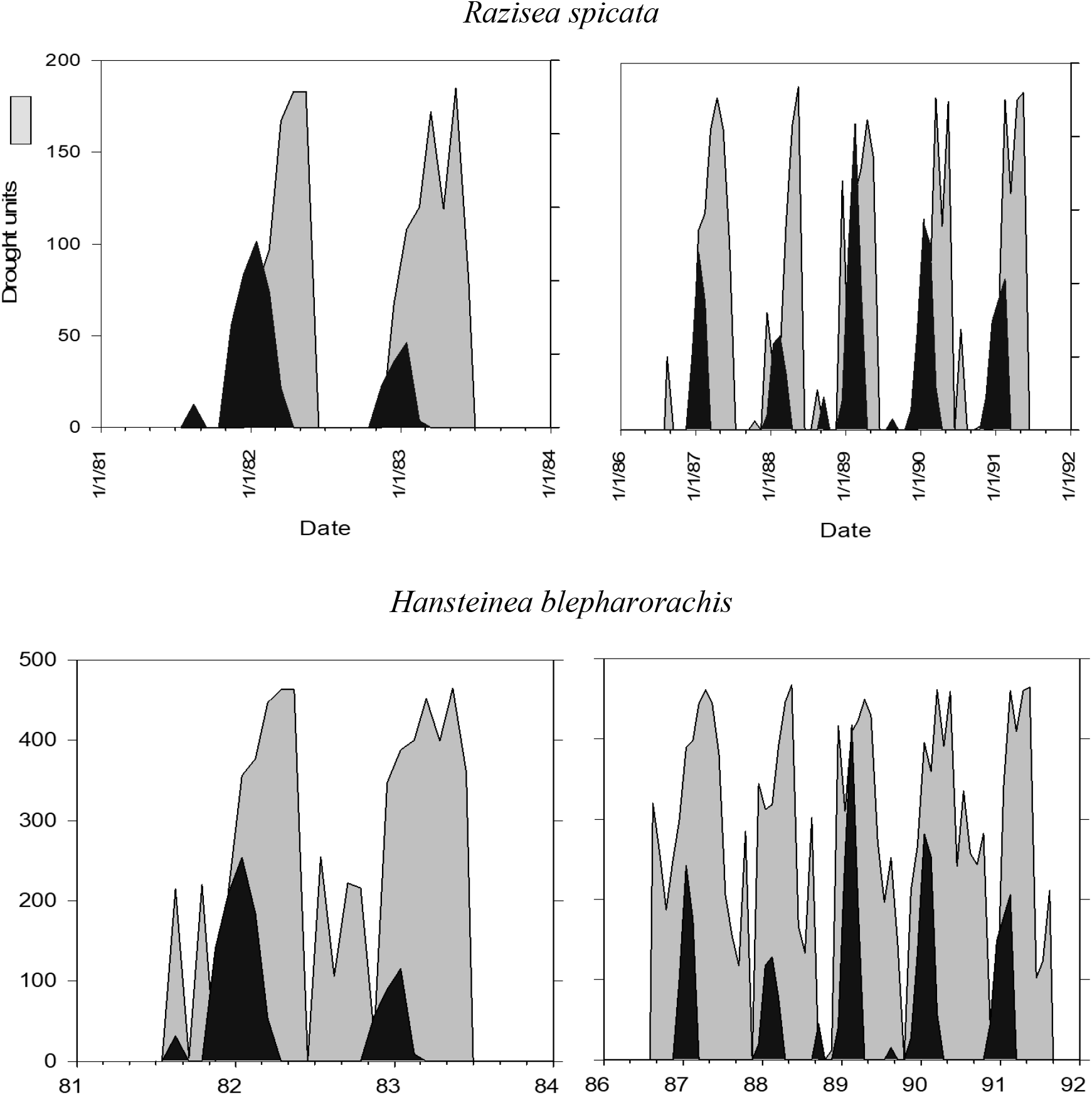
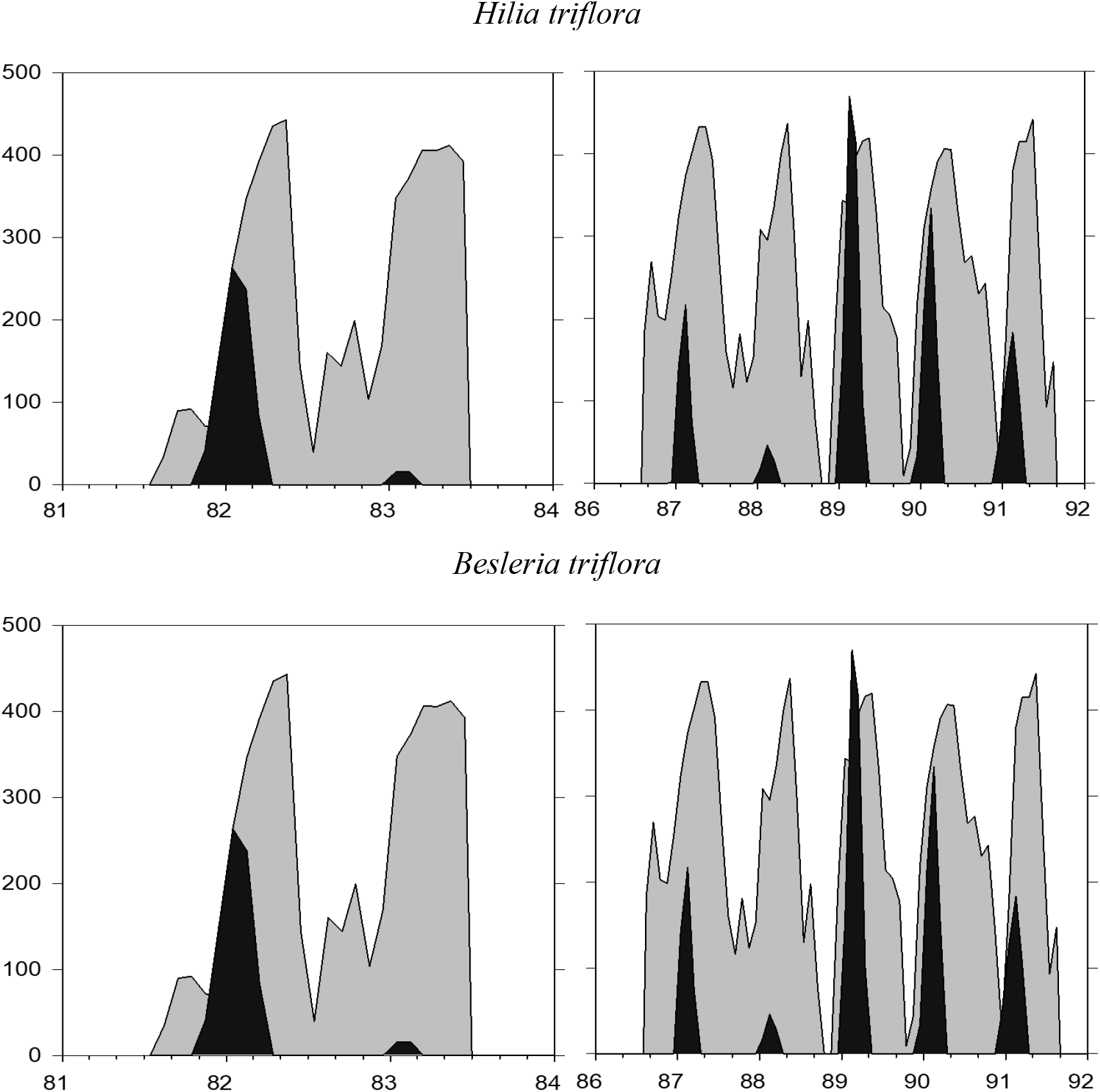
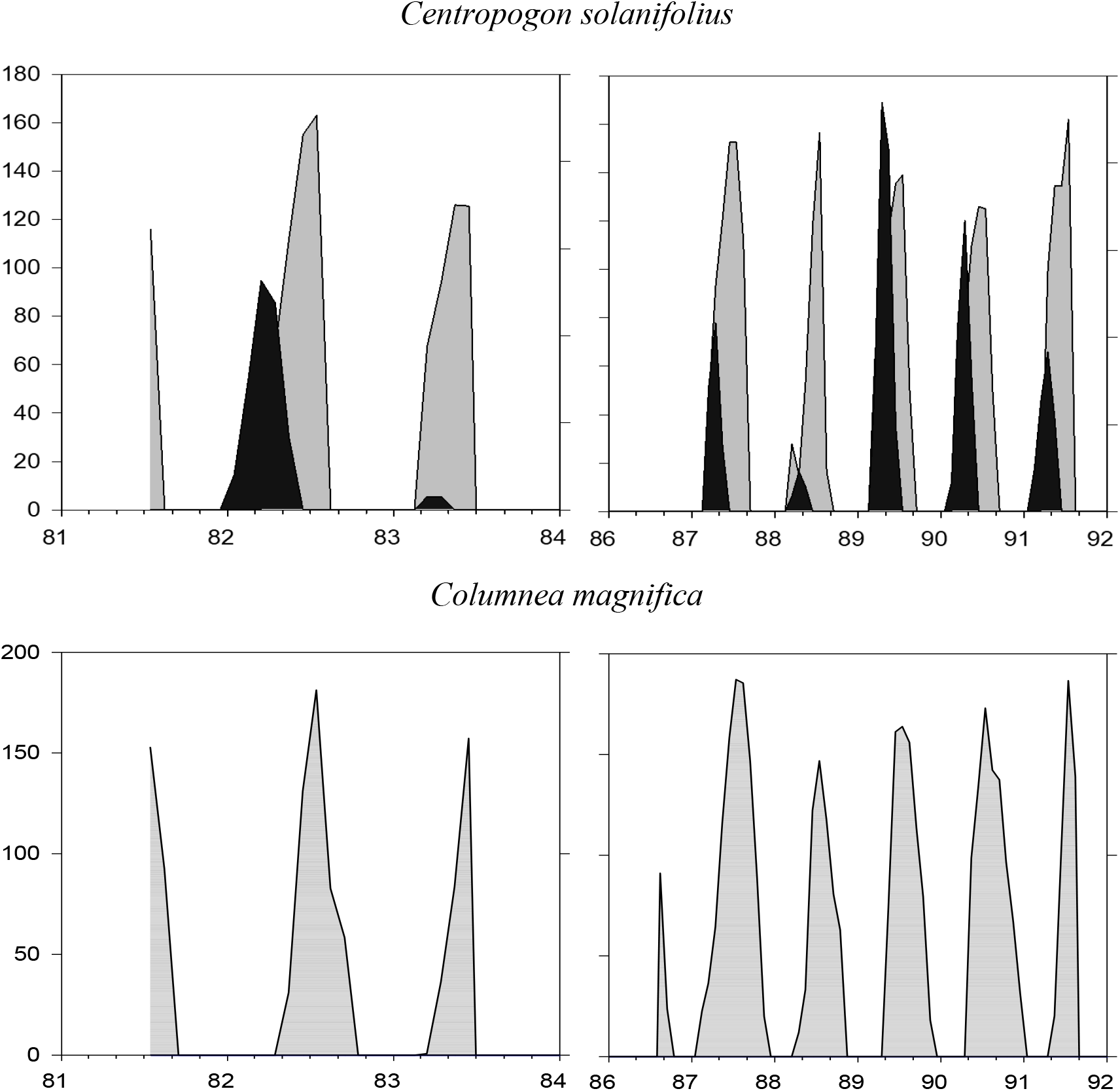
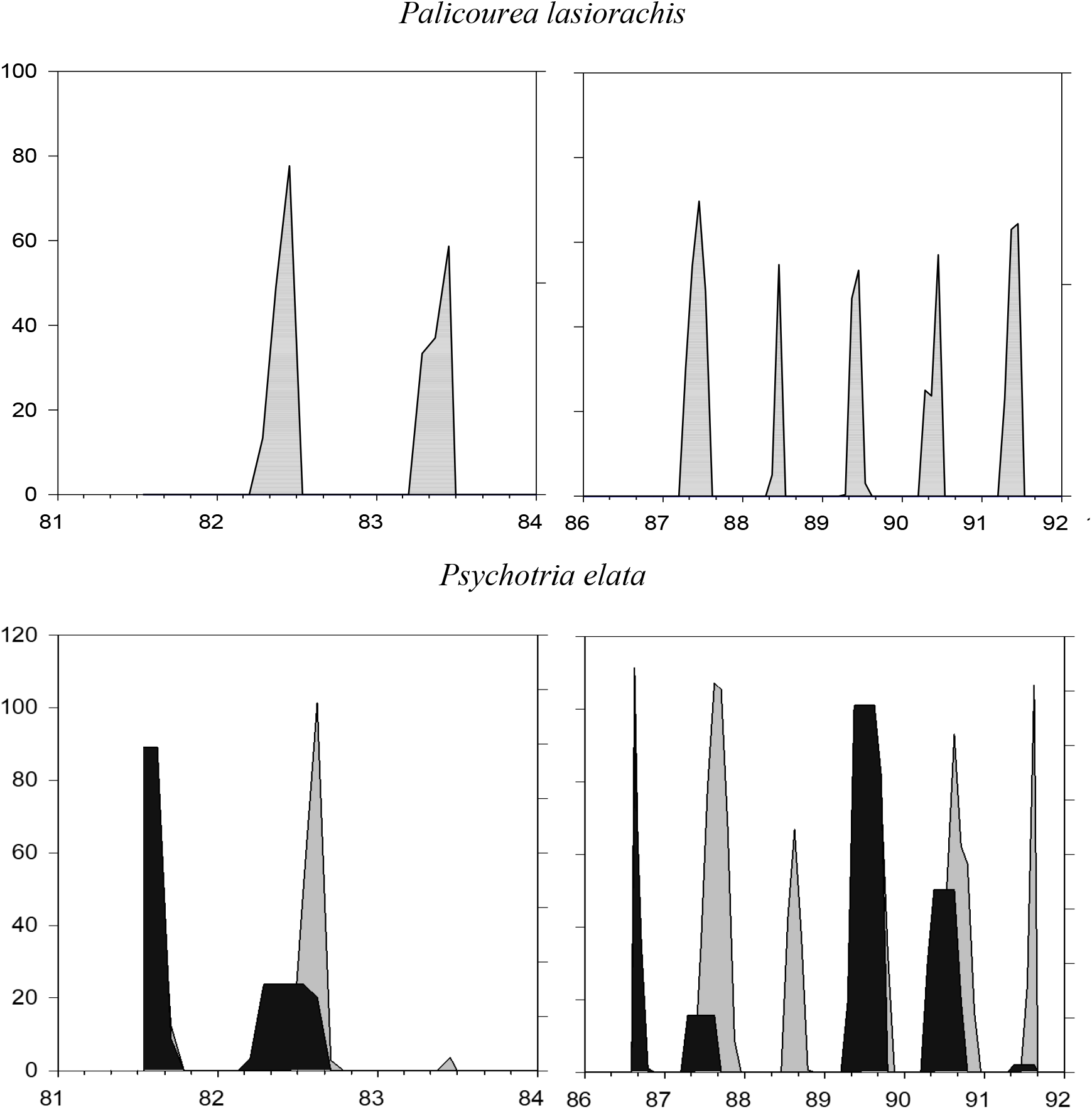
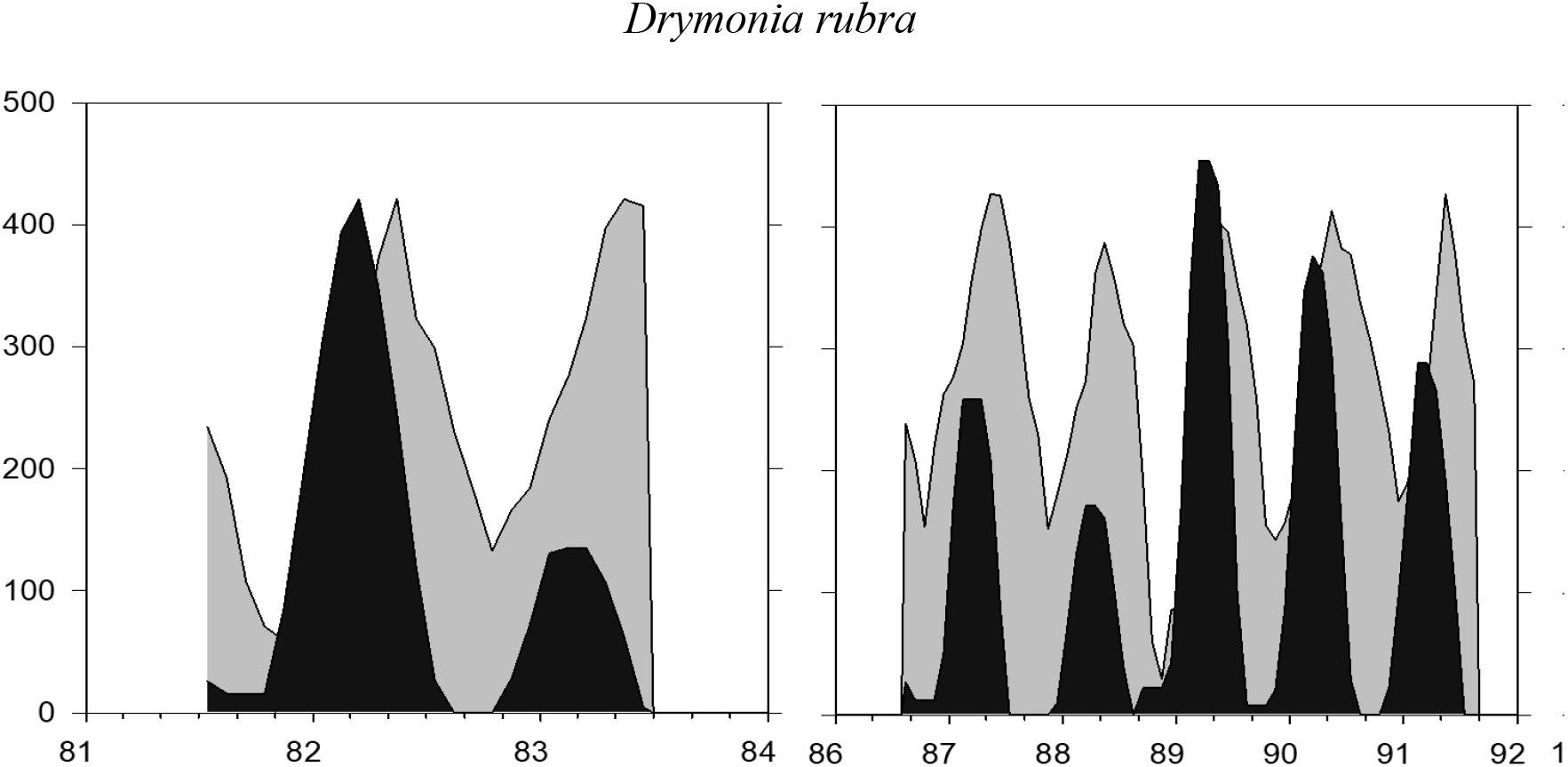
Estimated drought units (gray) and chill units (black) calculated for each species over time based on the optimal model output from model selection (see text). The three sampling periods are P1= July 1981-June 1983; P2= August 1986-August 1981; and P3=January 2017 to June 2018.

The differences in threshold drought levels for different species suggests that there are species-specific sensitivities to cues. These differences are potentially important if climate change ultimately leads to long term shifts in either drought or chill units accumulated over a given period of time. The implication is that the timing of some species might be much more likely to be disrupted by climate change than others.

### 4.2 Implications

As in the lowlands of Malaysia (Chen et al. 2018), the majority of the species studied here flower in relation to a combination of both seasonal cooling and accumulated dry periods (Table 2). These have changed to some extent since the early 1980s. Pounds et al. (1999) found that between 1972 - 1997 there was a long-term increasing trend in minimum daily temperatures, which affects the accumulation of chill units.

The question of climate change might affect the mutualistic interaction between the plant species in this study and the hummingbirds for which they are an important source of energy remains open. Additional studies of the timing of interactions between hummingbirds and these plants will be informative.

Climate change effects on tropical mutualisms such as that between ornithophilous plants and their hummingbird pollinators may well extend beyond simple asynchrony (Kharouba et al. 2018) due to shifts in phenology. Both the timing and magnitude of important tropical cues for flowering including precipitation and temperature are susceptible to change with increasing global temperatures. Ultimately, to understand the consequences for biological interactions, we need to understand how plants evolve in response to changing cues.

## Acknowledgements

The majority of the data analyzed in this work was collected by Peter Feinsinger, Willow Zuchowski, Greg Murray and Yan Linhardt. We thank them for gratiously providing acess to their data. We gratefully acknowledge the assistance of the staff of the Reserva Biológica Bosque Nuboso Monteverde, Centro Científico Tropical, SINAC and the Monteverde Institute who made this work possible.

